# A High-Throughput Platform for Assessing Single Fly Learning and Memory: Individual *Drosophila* Olfactory Conditioner (iDOC)

**DOI:** 10.1101/2024.12.18.629135

**Authors:** Antonio Ortega, Erik Tedre, El-Sayed Baz, Chang-Hui Tsao, Chien-Chun Chen, Sha Liu

## Abstract

Olfactory conditioning in *Drosophila melanogaster* is a widely used behavioral paradigm for uncovering the neural basis of learning and memory. The traditional T-maze assay, however, relies on group-level assessments and cannot examine individual flies’ learning and memory abilities. To address this limitation, we developed the Individual *Drosophila* Olfactory Conditioner (iDOC), an apparatus for single-fly conditioning and memory assessment. Using iDOC, we demonstrated aversive olfactory learning, short-term memory (STM), anesthesia-resistant memory (ARM), and protein synthesis-dependent long-term memory (LTM) in individual flies, consistent with previous experiments conducted using the T-maze. As an individual memory assessment method, iDOC can be integrated with other behavioral assays at the single animal level, such as sleep monitoring. Notably, we found that spaced training, which induces LTM, also promotes post-learning sleep, while massed training, which induces ARM, did not. iDOC thus provides a powerful platform for investigating learning and memory and its interplay with other behavioral processes at the individual animal level.

**Highlights:**

- Developed iDOC for high-throughput, single-fly aversive olfactory training and memory assessment
- Demonstrated aversive learning, STM, ARM, and LTM in individual flies using iDOC
- Spaced conditioning, and not massed conditioning, promotes sleep

## Introduction

Learning and memory are essential for animals to adapt and thrive in their environments. *Drosophila melanogaster* is no exception, demonstrating noticeable learning and memory capabilities^1^. Pioneering studies established the fruit fly as a powerful model organism for investigating these processes, with the T-maze apparatus providing a foundational tool for studying aversive olfactory conditioning and measuring memory at the population level. This T-maze olfactory assay has significantly advanced our understanding of the genetic, molecular, cellular, and circuit mechanisms underlying learning and memory^2–5^.

In the T-maze assay, groups of approximately 100 flies are conditioned for one minute to associate one of two odors, designated as the conditioned stimulus (CS), with 12 electric shocks serving as the unconditioned stimulus (US). During testing, flies are given a choice between two T-maze arms—one containing the shock-paired odor (CS+) and the other containing the unpaired odor (CS−)—and their memory is inferred from the proportion of flies avoiding the CS+. This single session of odor-electric shock association produces short-term memory (STM), which is barely detectable after 24 hours. However, repeating the one-minute training sessions induces distinct forms of long-lasting memory, depending on whether the sessions are “spaced” with rest intervals or “massed” without rest. Spaced training leads to protein synthesis-dependent long-term memory (LTM), while massed training results in anesthesia-resistant memory (ARM), which is protein-synthesis-independent^3^.

However, while memory acquisition and consolidation occur at the level of individual files, the T-maze evaluates learning and memory at the population level and lacks individual resolution. In addition, this limitation makes it challenging to study interactions between learning, memory, and other behaviors typically assessed at the individual level, such as feeding and sleep. To address these challenges, we present the Individual *Drosophila* Olfactory Conditioner (iDOC), a high-throughput platform for olfactory training and memory measurements in individual flies. iDOC enables assessing aversive olfactory learning and various forms of memory depending on the training paradigms, with results consistent with those obtained using the T-maze. Additionally, iDOC provides the flexibility to investigate interactions between memory and internal state variables, such as sleep, at the single-animal level. Detailed documentation for constructing and using iDOC, including hardware and software requirements, is available at https://idoc-docs.readthedocs.io.

## Results

### Design and Validation of iDOC for Aversive Olfactory Conditioning in Flies

The iDOC system consists of an odor delivery system, an electric shock delivery system, and a camera, all controlled by a computer. This setup can connect to up to 20 behavioral chambers for high-throughput experiments. Each chamber includes two lateral inflows for delivering two distinct odors (or one odor as the conditioned stimulus, CS, and air) into the chamber (Figure 1A). These odors mix in a central decision zone and vent through two central outflows. The chambers are enclosed with two glass slides, enabling video monitoring of flies’ movements from above and infrared illumination from below. To deliver electric shocks as the unconditioned stimulus (US) to the flies, the glass slides are coated with a transparent and conductive indium tin oxide (ITO) circuit, connected to the electric shock delivery system via two electrodes on either side of the chamber (Figure 1A). The transparency of the ITO coating allows video recording of fly behavior during experiments and facilitates the integration of optogenetic stimuli.

**Figure 1.**
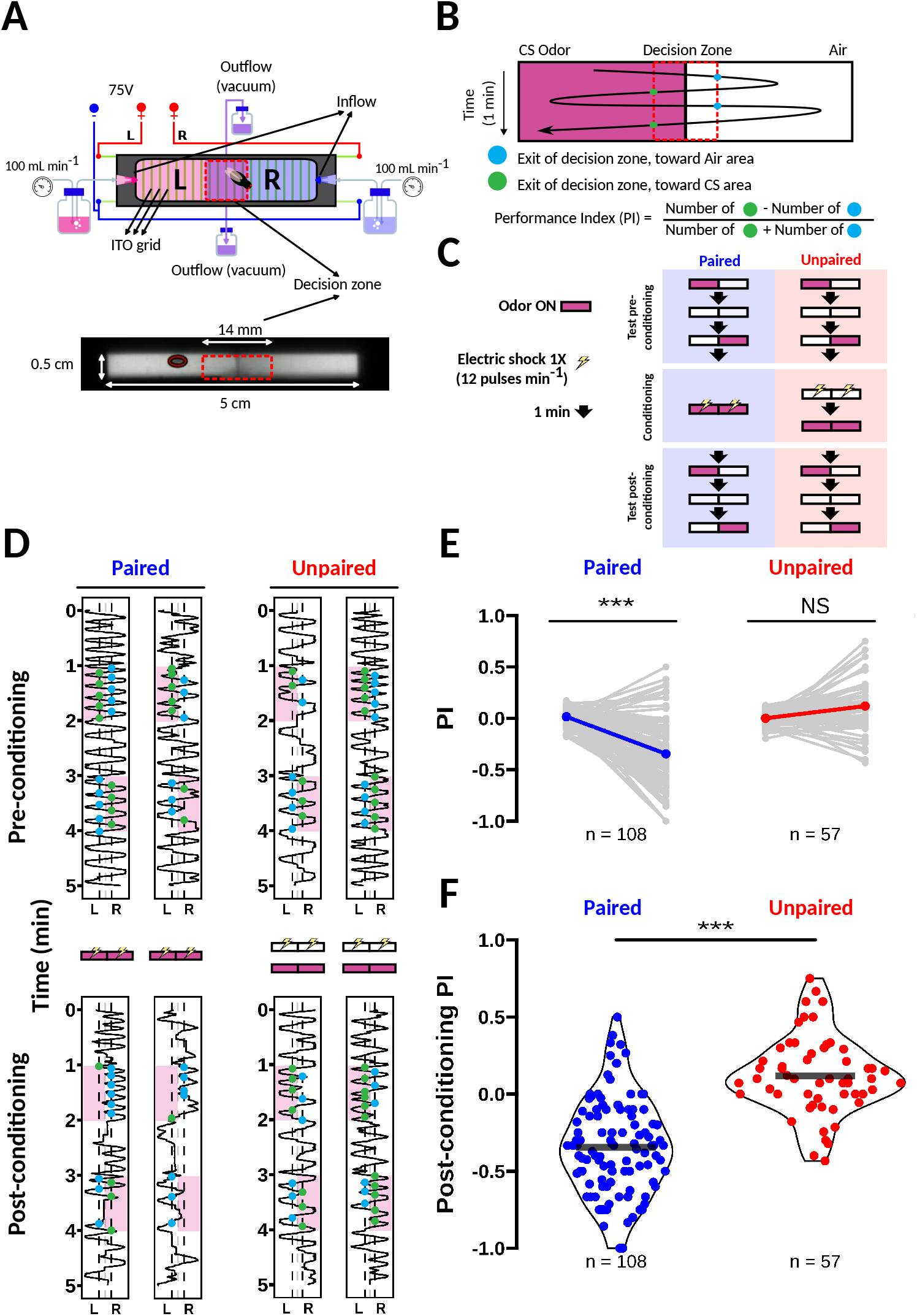
The iDOC system can form and recall memory in *Drosophila* using aversive olfactory conditioning. **A**. Schematic of the iDOC behavioral chamber, where the fly experiences all stimuli, including odor (Conditioned Stimulus or CS) delivered at 100 ml min^−1^ and 75V electric shock (Unconditioned Stimulus or US). **B**. The performance index, which assesses how strongly a fly avoids the CS, and therefore serves as a proxy of memory, is computed by counting exits from the decision zone and spans from -1 (aversion) to +1 (attraction). **C**. Experimental paradigm where US and CS are presented simultaneously (paired, left) or not (unpaired, right). The unpaired protocol serves as negative control for sensitization effects. **D**. Trajectory of two pairs of flies, undergoing the paired (left) and unpaired (right) protocols, each fly in its iDOC chamber, over time. Only the flies in the paired protocol acquire an aversion for the CS, represented by red rectangles, in the post-conditioning test. **E**. The average aversion acquired after conditioning in the paired paradigm is -0.35, while conditioning in the unpaired paradigm results in no acquired aversion. **F**. The two groups of flies therefore acquire a differential aversion for the CS. The horizontal black line represents the mean of each group.

A Python module tracks the flies’ real-time positions and delivers stimuli based on user-defined paradigms. This information is processed to detect exits from the decision zone and thus infer fly odor preference using a performance index (PI). The PI is calculated as the difference between exits toward the side with the conditioned stimulus (CS) and exits toward the side with air, divided by the total number of exits. The formula for the PI is:

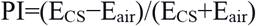

Here, E_CS_ represents the number of exits toward the side with the CS, and E_air_ represents the number of exits toward the side with air. The PI ranges from -1 to +1, where +1 indicates complete attraction to the CS (all exits toward the CS), 0 reflects neutrality (equal exits toward both sides), and -1 indicates complete aversion to the CS (all exits toward the side with air) (Figure 1B).

We first assessed the ability of iDOC to perform olfactory aversive learning by conditioning flies to avoid an odor paired with electric shocks. Olfactory aversive conditioning was induced by administering a cycle of 12 brief electric shock pulses (1 s duration at 0.2 Hz, 75 V) as the US on both sides of the chamber while simultaneously presenting the CS on both sides for 1 minute (Figure 1C). We measured the PI before and after training. During each test, the CS was presented on one side of the chamber without the US, and the test was repeated with the CS on the opposite side to control for side bias (Figure 1C). Flies exposed to the CS and US in a paired manner learned to avoid the CS (Figure 1D). In contrast, flies conditioned with an unpaired protocol, where the CS and US were not presented simultaneously, failed to develop an aversion to the CS after training (Figure 1D). On average, flies trained with the paired protocol achieved significant avoidance (PI<0), measured 20 minutes after training, while those subjected to the unpaired protocol showed no avoidance of the CS (Figure 1E). The significant difference in PI between the paired and unpaired groups confirms that aversive learning in flies requires the temporal co-occurrence of the CS and US, consistent with previous findings from the T-maze (Figure 1F). These findings demonstrate that iDOC is a reliable tool for olfactory aversive learning experiments in individual flies.

### Short-Term Memory Formation Following a Single Training Session

A single 1-minute conditioning session forms short-term memory (STM), which typically decays over a few hours, as found by previous T-maze-based experiments^3^. To validate that memory formed with one training session using iDOC follows similar dynamics, we measured the PI of flies at 20 minutes, 1 hour, 3 hours, and 24 hours after a single session of olfactory aversive conditioning (Figure 2A). As expected, memory formed by one session of training showed signs of decay after 1 hour and was completely absent after 24 hours (Figure 2B and 2C). These results confirm that the memory produced by a single short session of aversive olfactory conditioning is short lived, consistent with STM as previously described in the literature.

**Figure 2.**
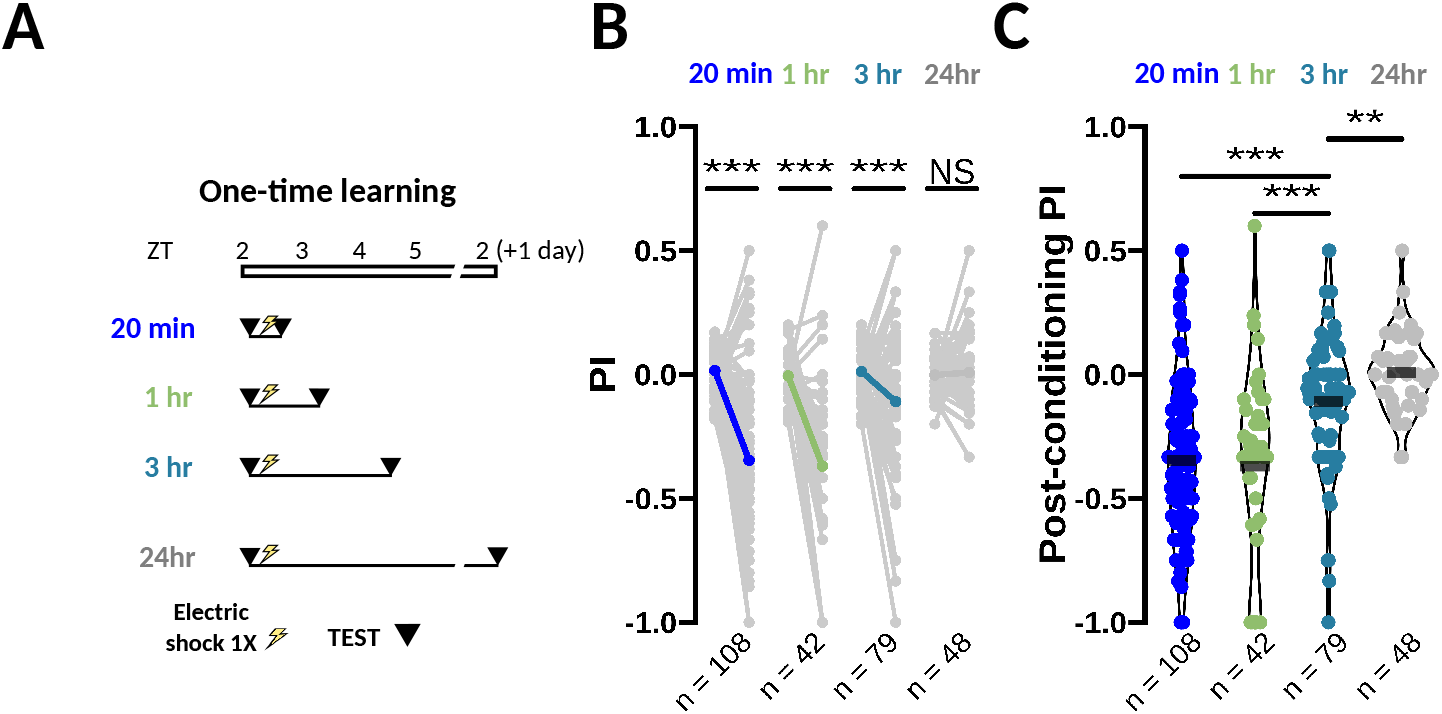
Design and Validation of iDOC for Aversive Olfactory Conditioning in Flies. **A** Experimental procedure followed to quantify memory persistence after one training round (12 pulses in 1 minute). The aversion for the CS is tested 20 minutes, 1 hour, 3 hours and 24 hours after conditioning. **B**. The acquired aversion is forgotten as the memory vanishes, reaching non-significance after 24 hours. **C**. The drop in aversion is significant already after 3 hours and follows a relatively linear trend toward neutrality for the CS after 24 hours.

### Protein Synthesis-Dependent Long-Term Memory Formation using iDOC

Unlike the single session of training used to induce short-term memory (STM) in flies, longterm memory (LTM) formation requires spaced training—multiple sessions separated by ∼15-minute rest intervals. This LTM persists for at least one day and can last a week or longer^3,6,7^. Importantly, LTM formation is protein synthesis-dependent, and disrupting the *de novo* protein synthesis during the consolidation phase prevents its formation^3^.

To evaluate LTM formation using iDOC, we implemented a spaced training paradigm consisting of six training sessions of the conditioned stimulus (CS) paired with the unconditioned stimulus (US), with 15–20-minute rest intervals between sessions (Figure 3A). This protocol mirrors the well-established spaced training paradigms used in T-maze experiments to induce robust LTM in *Drosophila*. Flies subjected to this paradigm formed LTM against the CS, which was recalled one day after training (Figures 3B and 3C). To confirm that this memory is protein synthesis-dependent, we pharmacologically inhibited protein synthesis through cycloheximide administration (CXM). Flies fed with CXM failed to form LTM, demonstrating that the memory induced by spaced training in iDOC requires active protein synthesis (Figures 3B and 3C). Additionally, we assessed flies carrying the *orb2*^*ΔQ*^ mutation, which disrupts the function of Orb2, a critical transcription factor in converting STM to LTM^8^. Consistent with previous findings with the T-maze apparatus, *orb2*^*ΔQ*^ mutants were unable to form LTM (Figures 3B and 3C). However, these mutants retained the ability to form STM, as evidenced by their significant avoidance of the CS 20 minutes after training (Figures 3B and 3C). These results validate the iDOC system as a reliable platform for inducing protein synthesis-dependent LTM in *Drosophila*.

**Figure 3.**
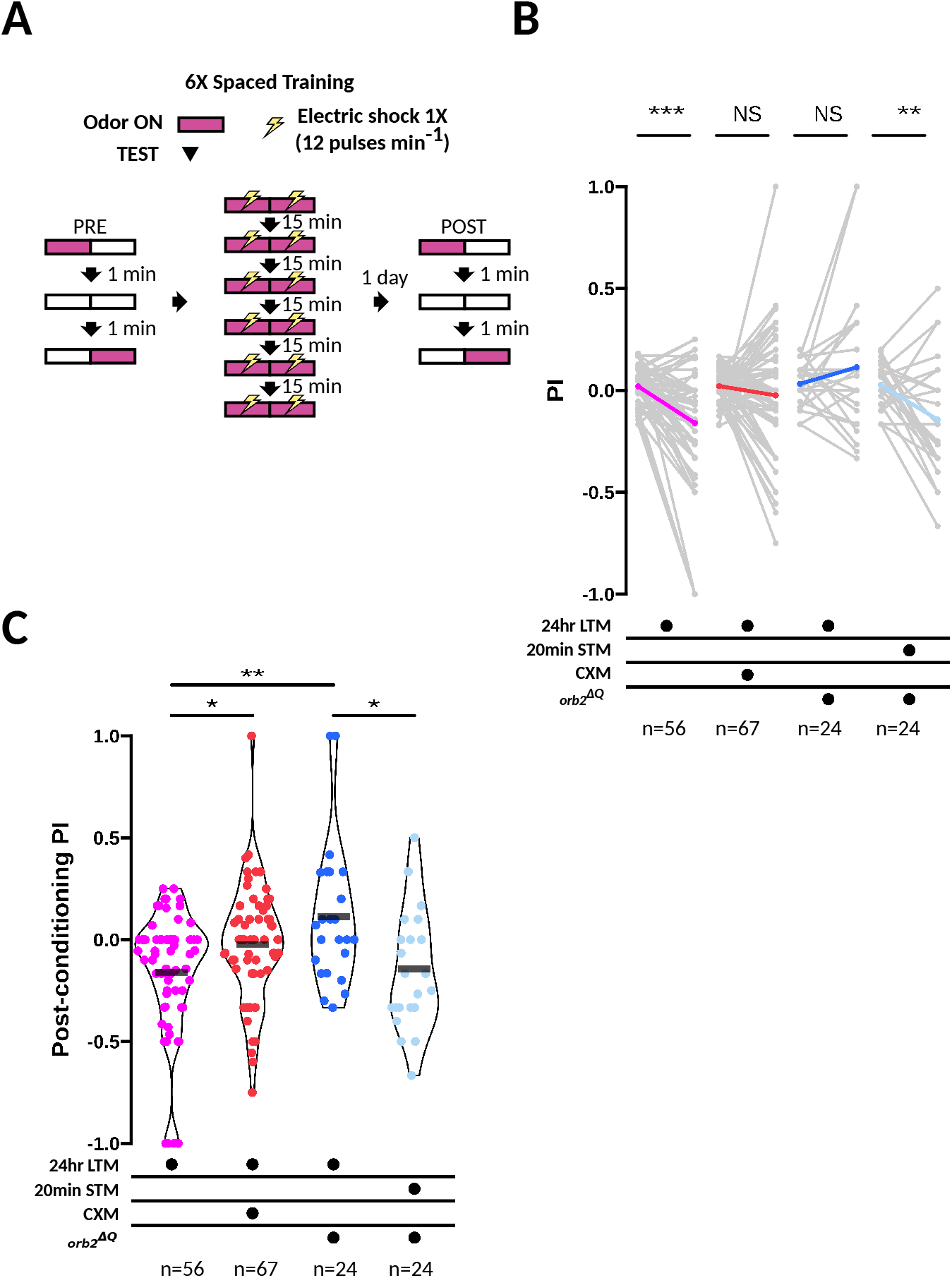
Protein Synthesis-Dependent Long-Term Memory Formation using iDOC. **A**. Experimental procedure used to generate protein-synthesis-dependent LTM. After running a pre-conditioning preference test, the animals undergo six training rounds of US and CS pairing, with a resting interval of 20 minutes between each round. The post-conditioning preference test is performed one day later. **B**. Pharmacological and genetic manipulations characterize the formed memory. After 24 hours, an aversion for the CS is still recalled in WT (wild type) animals (left). The administration of cycloheximide (CXM), a protein synthesis blocker, disrupts memory consolidation, resulting in a defective aversion after 24 hours (middle left). A deletion mutation in one of the transcription factors coordinating the required protein synthesis (Orb2) renders it non-functional and is followed by a loss of aversion after 24 hours (middle right). Notably, the learning ability of this mutant is not critically affected, suggesting a consolidation-specific problem (right). **C**. These manipulations reveal that the formed LTM lasts at least one day and is protein-synthesis dependent.

### Spaced Aversive Olfactory Conditioning Causes Increased Post-Learning Sleep

As a tool for assessing individual memory, iDOC can seamlessly integrate with other single-animal behavioral assays, such as sleep monitoring and manipulation. Studies in mammals and *Drosophila* suggest a strong role for sleep in memory consolidation^9–11^. In flies, long-term courtship conditioning has been shown to induce sleep after training^12^, while post-learning sleep is increased following appetitive olfactory conditioning when flies have access to food^13^. However, it remains unclear whether spaced aversive olfactory conditioning, which induces aversive olfactory LTM, also increases post-conditioning sleep.

To investigate whether and how sleep is modulated during the consolidation of protein synthesis long-term memory (LTM), we subjected flies to six sessions of spaced aversive olfactory conditioning. After 24 hours of sleep monitoring, we retrieved the flies and assessed the strength of their memory (Figure 4A). To control for the stimulus exposure (CS and US), we conditioned another group of flies using a massed training paradigm. In massed training, six sessions of aversive olfactory conditioning are administered consecutively without rest intervals, resulting in the formation of anesthesia-resistant memory (ARM)^14^, which is protein-synthesis-independent^3^ (Figure 4B). Sleep was quantified by transferring the animals into a sleep monitoring device that measures sleep duration while preserving individual fly identity (Figure 4C). Although both spaced and massed training paradigms produced memories that could be retrieved 24 hours after training (Figure 4D), only flies subjected to the spaced paradigm exhibited an increase in sleep during the hours immediately following conditioning, specifically between *zeitgeber* (hours since lights on, ZT) 5 and 11 (Figure 4E). In contrast, sleep amounts were indistinguishable across spaced, massed, and untrained control groups between ZT12 and ZT18 (Figure 4F). These findings indicate that protein-synthesis-dependent LTM induces increased sleep during the early post-training phase, likely reflecting a critical window for memory consolidation. Taken together, these results highlight how iDOC can be used to study the complex interplay between memory consolidation and sleep in individual flies, enabling high-throughput investigation of these processes.

**Figure 4.**
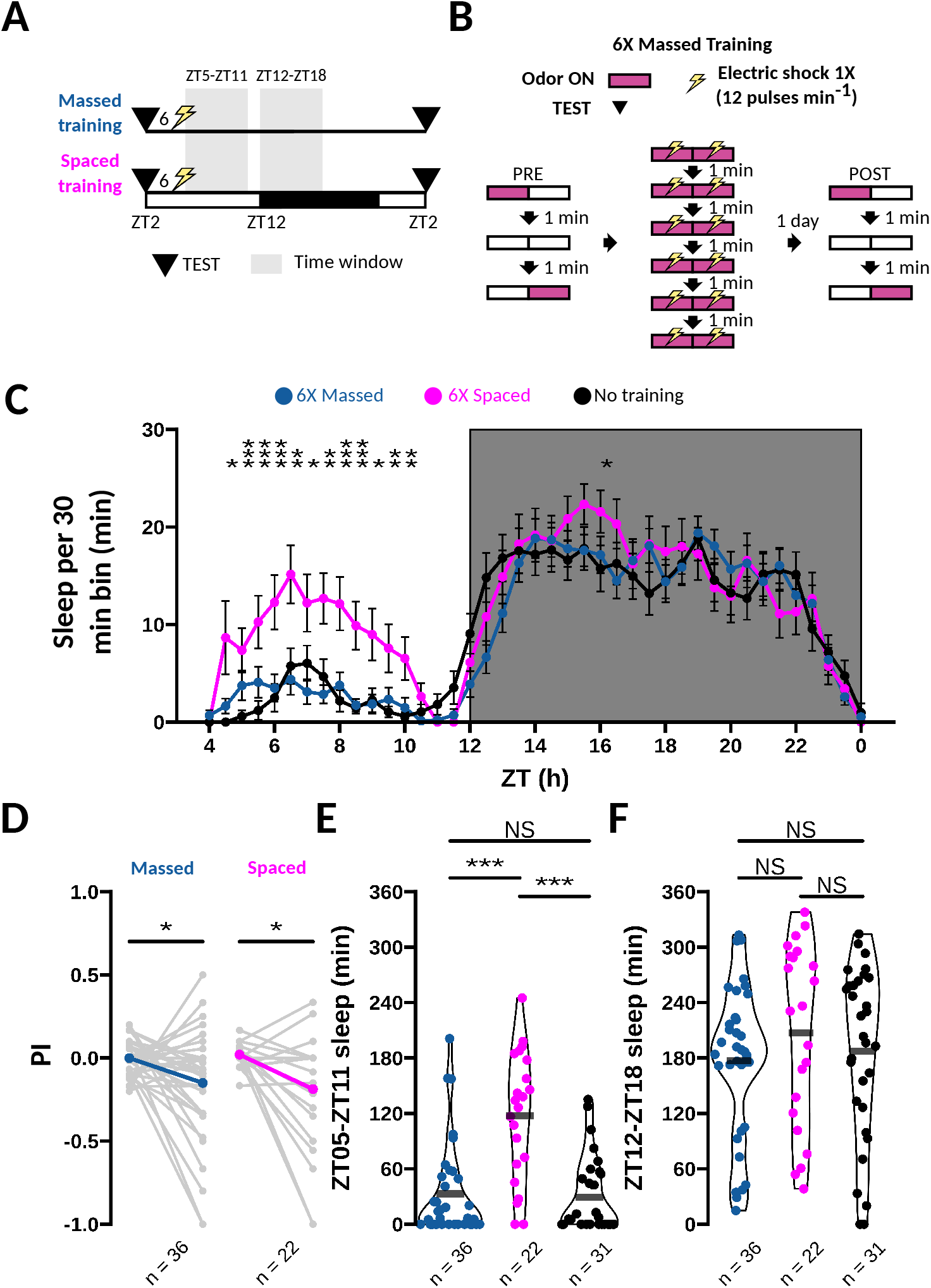
Spaced Aversive Olfactory Conditioning Causes Increased Post-Learning Sleep. **A**. Experimental procedure used to generate protein-synthesis-independent long-term memory (anesthesia-resistant memory, ARM). Like in spaced training, the animals undergo six rounds of conditioning. These however are only interspersed by oneminute intervals. **B**. During the consolidation process, sleep is monitored in separate tubes, until the test one day after conditioning. **C**. Sleep is quantified by counting every 30 minutes the time the animal spends in the sleep state using the Ethoscope. **D** Both massed and spaced protocols create an aversion toward the CS 24 hours after conditioning. **E**. An increased amount of sleep during the 6 hours after conditioning (ZT5-ZT11) was observed in the flies which experienced spaced training (middle), as compared to those which underwent massed training (left) or no training at all (right). **F**. This difference subsides in the following six hours (ZT12-ZT18), with all groups sleeping a similar amount.

## Discussion

Here, we introduced iDOC, an apparatus designed for aversive olfactory conditioning, capable of assessing both short- and long-term memory in individual flies. Unlike the T-maze-based assay, which provides a single group-level memory readout per experiment, iDOC enables the simultaneous assessment of up to 20 individual flies, significantly increasing experimental throughput. By replicating well-established associative learning paradigms established in T-maze, our results validate iDOC as a reliable and versatile platform for studying learning and memory in *Drosophila*. To facilitate its adoption, we provide comprehensive documentation on how to build and operate iDOC, as well as detailed protocols for data analysis (https://idoc-docs.readthedocs.io).

Our findings using iDOC reveal that spaced training induces sleep during a critical six-hour window (ZT5-ZT11). This observation implies that sleep deprivation during this period could disrupt memory consolidation. Consequently, we hypothesize that flies subjected to the spaced training paradigm but prevented from sleeping will display a more neutral distribution of performance indices, reflecting impaired memory consolidation.

Two other apparatuses for conditioning and assessing olfactory memory in individual flies have been previously reported. The pioneering apparatus developed by Claridge-Chang *et al*.^*15*^ was the first to enable aversive olfactory conditioning in individual flies. However, it relied on non-transparent copper floor and ceiling circuit boards to deliver electric shocks, requiring side illumination for video recording. This design introduced positional bias in the light stimulus strength based on the fly's location within the chamber when combined with optogenetic stimulations. Therefore, the apparatus could only accommodate four chambers when combined with optogenetic stimulation, significantly limiting experimental throughput. A more recent apparatus, published while we finished our experiments^16^, introduced the usage of indium tin oxide (ITO)-coated circuit boards to deliver electric shock US, similar to the iDOC design. In any case, our apparatus is, to the best of our knowledge, the first one to demonstrate LTM formation in individual flies.

In addition to higher throughput, measuring learning and memory in individual flies offers several key advantages over group-based assays. First, individual assays eliminate the confounding effects of group interactions. While the original T-maze experiments measures the performance from population of flies^4^, a recent study has shown that the behavior of naïve flies can be influenced by mechanosensory interactions within the group, potentially introducing bias or variance as subsets of the population influence each other's actions^17^. Isolating flies during experiments fully prevents such group effects, ensuring that behavioral responses are not confounded by social interactions.

Second, group-based assays, such as the T-maze, often require flies to choose between two odors: one coupled to the US (CS+) and another uncoupled (CS-). While this design ensures a balanced 50:50 distribution in naïve flies, it introduces an undesired "safety memory" effect driven by the CS-, rather than the CS+^18^. By omitting the CS-, iDOC simplifies the experimental design and avoids these confounding effects, ensuring that the memory readout is solely based on only one CS. It also reduces the complexity of studying the combined memory processes in the brain.

Finally, individual memory assessment in iDOC facilitates the maintenance of animal identity, enabling the correlation of internal state variables, such as sleep, with learning and memory performance. In a complementary study conducted in our laboratory, we paired learning scores from iDOC with prior sleep behavior, revealing how sleep and wakefulness influence learning ability (Chen *et al*., in preparation). Future studies could leverage iDOC to investigate the effects of manipulating other internal state variables, such as hunger or thirst, on learning and LTM formation, providing new insights into the interplay between physiological states and memory.

### Limitations of the Study

A consideration for iDOC is that the precision of the memory readout (PI) relies on the animal's activity levels. Low locomotor activity can prevent sufficient exploration to distinguish preference from chance, making memory assessment less reliable. To address this, we conducted experiments shortly after the dark-light transition (ZT1–ZT4), when flies are typically more active, though conducting experiments before the light-dark transition is also a viable option. Alternatively, using the time spent in the CS area as a proxy for memory could mitigate the impact of low activity levels, as this measure is less dependent on locomotion than exits from the decision zone.

## STAR Methods

### Key resources table

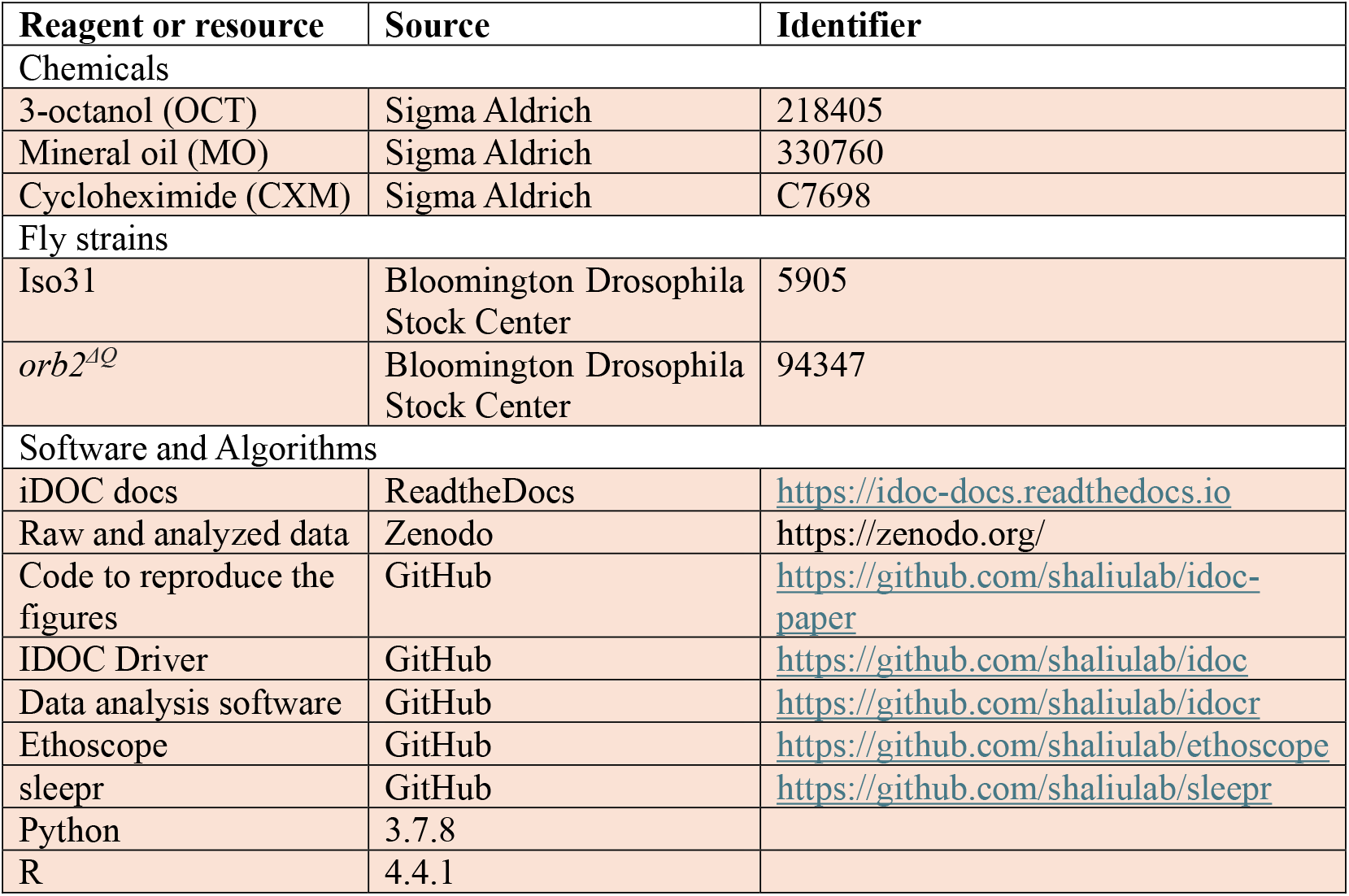

### Experimental model and subject details

#### Fly

4-7 days old mated female *Drosophila melanogaster*, entrained in a 12:12 h light:dark cycle since eclosion on a standard cornmeal-agar diet, were used for all experiments. The flies were kept in small vials of 10-20 females with 3-5 males, at 22°C and approximately 70% humidity during entrainment. Female flies were preferred due to their increased size, which improved the efficiency of electric shock delivery. Experiments were conducted from ZT1 to ZT4, which is when the animals exhibit their peak activity.

#### Preparation of odors

The sources of odor were prepared by loading clean mineral odor and a fixed volume of the odorant into a single bottle and mixing both reactants by bubbling pressurized air for three minutes. Acuity tests showed that neutral preference was optimized when OCT was diluted with a factor of 1:500, or 0.2% (data not shown). This dilution was achieved by loading 120 μl of OCT into 60 ml of MO. The homogenized mix was split into two bottles of 30 ml. MO bottles were made by loading 30 ml of mineral oil into two different bottles.

#### Pavlovian aversive conditioning

12 rounds of 1 second electric shock paired with odor were delivered to each chamber to couple CS and US. The CS was provided by both odor inflows on either side of the chamber simultaneously, with a flowrate of 100 ml min^-1^. Electric shocks were delivered through a conductive grid of indium tin oxide (ITO) laid on the glass slides enclosing the chambers, using a 75V power source.

#### Assessment of memory

Avoidance behavior against the CS was assessed as a proxy for memory strength. This avoidance was quantified before and after conditioning by delivering the CS during two separate 1-minute trials on alternating sides of the chamber, while delivering neutral MO odor on the other side. The swapping of CS and MO in the second trial helped control for side-bias effects.

The delivery of CS and MO on each side of the chamber generates a mixing area or decision zone around the center of the chamber. The decision zone was set to spread 7 mm in either direction from the center, based on observations of fly behavior. Exits out of this zone toward either side can be used as proxy of preference for the stimulus on the exiting side. Therefore, one can use these exits to build a statistic that quantifies the preference for the CS. Specifically, a performance index (PI) can be computed by counting the exits toward the MO side, subtracting the counts toward the CS side, and dividing this difference by the sum of exits This yields a number between -1 and 1, where -1 represents pure avoidance or aversion, and +1 pure attraction. Alternatively, the fraction of time spent on the CS side could be used but it would be biased by immobility bouts which are independent of stimulus preference. The coordinates of the middle of the chamber were manually annotated using the calibration tool in this repository https://github.com/shaliulab/midline-detector.

#### Long-term memory experiments

Fly identity during the 24-hour period between conditioning and the post-training test was maintained by transferring each individual fly to a locomotor tube and storing the identity mapping. Protein-synthesis-dependent memory was disturbed in WT animals by providing 35 mM cycloheximide in the food for one day before the conditioning experiment and during the ensuing consolidation process.

#### Sleep quantification algorithm

All sleep recordings were performed in glass tubes loaded into an ethoscope^19^ which records the x, y coordinates of up to 20 animals per batch at a framerate of 2 FPS. Offline scoring of the behavior was performed in R using the following algorithm for each animal independently: First, the x,y time series were split in windows of 10 seconds each. Then, the algorithm tests whether the distance crossed by the animal's centroid between two neighboring time points (i.e. within 0.5 seconds) is greater than a threshold of 0.3 mm. If any pair of consecutive points implies a displacement greater than this distance, the fly is set to be moving for the whole 10 second window, otherwise it is not moving. The fly is set to be asleep if at least 30 10-second windows (i.e. 300 seconds or 5 minutes) in a row are spent in the non-moving state. The number of minutes spent in the sleep state is computed every 30 minutes as the number of 10-second windows that the animal spent in the sleep state, multiplied by 10 seconds. This algorithm is described in the sleepr package from the rethomics suite for analysis of ethoscope data^20^.

## Data analysis

The average of the PI observed on either trial was used as a representative metric of the memory demonstrated by the animal. If one of the trials did not feature three exits or more, only the other trial was used. Flies were discarded if 1) no trials had at least 3 exits 2) if a preference for one side, independent of the stimulus presented on that side, was demonstrated by the animal or 3) if a pre-conditioning bias for or against the odor was measured (absolute value of PI being greater than 0.2). Side-preference was manually scored. Finally, flies with a behavioral trace that clearly shows avoidance, but which doesn't strictly fit to the rules explained above were also manually scored.

A significant difference between pre-training and post-training tests was assessed using the one-sided paired Wilcoxon signed-rank test implementation in R's stats package (`wilcox.text`). Significant differences in PI scores achieved by flies in different groups were assessed with the one-sided unpaired Wilcoxon signed-rank test implementation. Multiple testing correction was performed using the Bonferroni method, by dividing the significance values or alpha (0.05, 0.01 and 0.005) by the number of tests performed. P-values are thus represented with one, two or three asterisks (*) depending on whether they lie below the first, second or third updated alpha value.

## Data availability

All raw data from all recordings used in this paper were uploaded to Zenodo.

## Code availability

All the R code required to generate the figures in the paper is available at https://github.com/shaliulab/idoc-paper.

## References

1. Quinn, W.G., Harris, W.A., and Benzer, S. (1974). Conditioned Behavior in Drosophila melanogaster. Proceedings of the National Academy of Sciences 71, 708–712. 10.1073/PNAS.71.3.708.

2. Dudai, Y., Jan, Y.N., Byers, D., Quinn, W.G., and Benzer, S. (1976). dunce, a mutant of Drosophila deficient in learning. Proceedings of the National Academy of Sciences 73, 1684–1688. 10.1073/PNAS.73.5.1684.

3. Tully, T., Preat, T., Boynton, S.C., and Del Vecchio, M. (1994). Genetic dissection of consolidated memory in Drosophila. Cell 79, 35–47. 10.1016/0092-8674(94)90398-0.

4. Tully, T., and Quinn, W.G. (1985). Classical conditioning and retention in normal and mutant Drosophila melanogaster. Journal of Comparative Physiology A 157, 263–277. 10.1007/BF01350033/METRICS.

5. Tully, T. (1984). Drosophila Learning: Behavior and biochemistry. Behav Genet 14, 527–557. 10.1007/BF01065446/METRICS.

6. McGaugh, J.L. (1966). Time-Dependent Processes in Memory Storage. Science (1979) 153, 1351–1358. 10.1126/SCIENCE.153.3742.1351.

7. Carew, T.J., Pinsker, H.M., and Kandel, E.R. (1972). Long-term habituation of a defensive withdrawal reflex in aplysia. Science 175, 451–454. 10.1126/SCIENCE.175.4020.451.

8. Keleman, K., Krüttner, S., Alenius, M., and Dickson, B.J. (2007). Function of the Drosophila CPEB protein Orb2 in long-term courtship memory. Nature Neuroscience 2007 10:12 10, 1587–1593. 10.1038/nn1996.

9. Le Glou, E., Seugnet, L., Shaw, P.J., Preat, T., and Goguel, V. (2012). Circadian Modulation of Consolidated Memory Retrieval Following Sleep Deprivation in Drosophila. Sleep 35, 1377–1384. 10.5665/SLEEP.2118.

10. Graves, L.A., Heller, E.A., Pack, A.I., and Abel, T. (2003). Sleep Deprivation Selectively Impairs Memory Consolidation for Contextual Fear Conditioning. Learning & Memory 10, 168–176. 10.1101/LM.48803.

11. Vecsey, C.G., Baillie, G.S., Jaganath, D., Havekes, R., Daniels, A., Wimmer, M., Huang, T., Brown, K.M., Li, X.Y., Descalzi, G., et al. (2009). Sleep deprivation mimpairs cAMP signalling in the hippocampus. Nature 2009 461:7267 461, 1122–1125. 10.1038/nature08488.

12. Ganguly-Fitzgerald, I., Donlea, J., and Shaw, P.J. (2006). Waking experience affects sleep need in Drosophila. Science (1979) 313, 1775–1781. 10.1126/SCIENCE.1130408/SUPPL_FILE/GANGULY.SOM.PDF.

13. Chouhan, N.S., Griffith, L.C., Haynes, P., and Sehgal, A. (2020). Availability of food determines the need for sleep in memory consolidation. Nature 2020 589:7843 589, 582–585. 10.1038/s41586-020-2997-y.

14. Quinn, W.G., and Dudai, Y. (1976). Memory phases in Drosophila. Nature 1976 262:5569 262, 576–577. 10.1038/262576a0.

15. Claridge-Chang, A., Roorda, R.D., Vrontou, E., Sjulson, L., Li, H., Hirsh, J., and Miesenböck, G. (2009). Writing Memories with Light-Addressable Reinforcement Circuitry. Cell 139, 405–415. 10.1016/J.CELL.2009.08.034/ATTACHMENT/F2588B1A-0FF0-4D78-A1F9-027C48D90C8A/MMC4.ZIP.

16. Matthew A.-Y. Smith, K.S.H.G.T. and B. de B.P.F. 2022 https://doi.org/10.1098/rsbl.2021.0424 (2022). Idiosyncratic learning performance in flies. Biol Lett 18. 10.1098/rsbl.2021.0424.

17. Ramdya, P., Lichocki, P., Cruchet, S., Frisch, L., Tse, W., Floreano, D., and Benton, R. (2014). Mechanosensory interactions drive collective behaviour in Drosophila. Nature 2014 519:7542 519, 233–236. 10.1038/nature14024.

18. Jacob, P.F., and Waddell, S. (2020). Spaced Training Forms Complementary Long-Term Memories of Opposite Valence in Drosophila. Neuron 106, 977–991.e4. 10.1016/J.NEURON.2020.03.013/ATTACHMENT/4BE2BFF7-6695-4B7B-A754-057525CC1913/MMC2.PDF.

19. Geissmann, Q., Garcia Rodriguez, L., Beckwith, E.J., French, A.S., Jamasb, A.R., and Gilestro, G.F. (2017). Ethoscopes: An open platform for high-throughput ethomics. PLoS Biol 15, e2003026. 10.1371/JOURNAL.PBIO.2003026.

20. Geissmann, Q., Rodriguez, L.G., Beckwith, E.J., and Gilestro, G.F. (2019). Rethomics: An R framework to analyse high-throughput behavioural data. PLoS One 14, e0209331. 10.1371/JOURNAL.PONE.0209331.

